# Extinction rates in tumor public goods games

**DOI:** 10.1101/134361

**Authors:** Philip Gerlee, Philipp M. Altrock

**Affiliations:** Department of Mathematical Sciences, Chalmers University of Technology, SE-41296 Gothenburg, Sweden; Department of Mathematical Sciences, University of Gothenburg, SE-40530 Gothenburg, Sweden; Department of Integrated Mathematical Oncology, Moffitt Cancer Center and Research Institute, Tampa, FL 33612, USA; University of South Florida Morsani College of Medicine, Tampa, FL 33612, USA

## Abstract

Cancer evolution and progression are shaped by cellular interactions and Darwinian selection. Evolutionary game theory incorporates both of these principles, and has been proposed as a framework to understand tumor cell population dynamics. A cornerstone of evolutionary dynamics is the replicator equation, which describes changes in the relative abundance of different cell types, and is able to predict evolutionary equilibria. Typically, the replicator equation focuses on differences in relative fitness. We here show that this framework might not be sufficient under all circumstances, as it neglects important aspects of population growth. Standard replicator dynamics might miss critical differences in the time it takes to reach an equilibrium, as this time also depends on cellular turnover in growing but bounded populations. As the system reaches a stable manifold, the time to reach equilibrium depends on cellular death and birth rates. These rates shape the timescales, in particular in co-evolutionary dynamics of growth factor producers and free-riders. Replicator dynamics might be an appropriate framework only when birth and death rates are of similar magnitude. Otherwise, population growth effects cannot be neglected when predicting the time to reach an equilibrium, and cell type specific rates have to be accounted for explicitly.

## 1 Introduction

The theory of games was devised by von Neumann and Morgenstern [1], and according to Aumann [2], game theory is an “interactive decision theory”, where an agent’s best strategy depends on her expectations on the actions chosen by other agents, and vice versa. As a result, “the outcomes in question might have been intended by none of the agents” [3]. In order to rank and order strategies, and to optimize individual payoffs, different systems to systematically identify equilibria have been defined. Most famously, the Nash equilibrium is a set of strategies such that no single agent can improve by switching to another strategy [4]. This concept includes mixed equilibria, which describe probability distributions over strategies. Such equilibrium concepts in game theory cover various kinds of patterns of play, i.e. simultaneous, non-simultaneous, and asymmetric strategies [5]. This rich and complex framework allows for a wide application of game theory beyond economics, famously in ecology and evolution [6]. In biological context, and especially in evolutionary game theory, the focus has been on simultaneous and symmetric strategic interactions in evolving populations [7].

Evolutionary game theory replaces the idea of choice and rationality by concepts of reproduction and selection in a population of evolving individuals [8] and was conceived to study animal conflict [9]. Behavioral phenotypes are hardwired to heritable genotypes. Without the possibility of spontaneous mutation events, offspring carry the parent strategy. Evolutionary games have also been used extensively to study learning and pairwise comparison-based changes in strategy abundance in populations of potentially erroneous players [10, 11, 12].

Selection in evolutionary games is based on the assumption that payoff translates into Darwinian fitness, which is a measure for an individual’s contribution to the pool of offspring in the future. Complex deterministic dynamical systems arise when one considers very large populations of reproducing individuals. The most prominent example for such a system is the replicator equation [13], which focuses on the relative abundance of each strategy. The replicator equation does not model population growth specifically, but rather describes changes in relative abundances. Existence and stability of fixed points in these dynamical systems depend on the payoffs [14], and on the choice of fitness function [15]. In the study of animal behavior, the precise measurements of payoffs, as observed from individuals’ behaviors, is difficult. Milinski *et al.* determined all but one payoff parameter precisely, in order to observe tit-for-tat strategies in repeated Prisoner’s Dilemma games in fish [16]. Kerr *et al.* showed that *E. coli* bacteria can be observed to evolve according to rock-paper-scissors type of interactions, if cellular dispersal is minimal As seen by a recent expansion of interesting theoretical considerations focusing on evolutionary games in biology [17] lies in the ability to assess many problems in ecological and evolutionary population dynamics at least in qualitative terms, i.e. by predicting and ranking evolutionary equilibria, how population-wide coexistence can emerge from apparent individual conflict, or how fast transitions between equilibria occur.

Tumor cell populations, including cells of the tumor microenvironment, are part of a complex ecosystem [18], which can have consequences for therapeutic outcomes [19]. At the same time, it has been more widely recognized that Darwinian selection plays a key role in cancer [20]. Given the appreciated amount of both genetic and phenotypic heterogeneity in tumor cell populations [21], evolutionary games have become more widely used as a means to theoretically model tumor evolution, especially after tumor initiation [22]. Prominent examples of recent applications of replicator equations in cancer are concerned with the avoidance of the tragedy of the commons, where a sub-population of tumor cells produces a ‘tumor public good’ in form of an insulin-like growth factor [23], in form of glycolytic acid and vascular endothelial growth factor [24], or modeling the dynamic equilibrium between lactate respiration and glycolysis in tumor cells [25]. Such non-autonomous effects between tumor cells had been proposed some time ago [26], and non-cell-autonomous growth rates were recently measured empirically [27]. Similar findings and future challenges in this field have been summarized by Tabassum and Polyak [18].

We here focus on the time it takes to reach an equilibrium in different approaches to model deterministic evolutionary game dynamics. In particular we focus on differences between logistic growth and the replicator dynamics. We show that the time to get arbitrarily close to an equilibrium, which we here call the *ε*-fixation time, might critically depend on the underlying cellular birth and death rates. We focus on two co-evolving tumor cell populations, and present a discussion of the dynamics between growth factor producers *C*_1_ and free-riding non-producers *C*_2_. In the simplest setting we can assume that these closely related cell types experience population doubling rate *α*, and that the tumor public good, produced by *C*_1_ cells, has a linear positive effect on cellular birth rates in form the additive benefit that scales with the doubling rate *βα*, but bears a production cost *κ*. The respective game can be recast in the payoff matrix

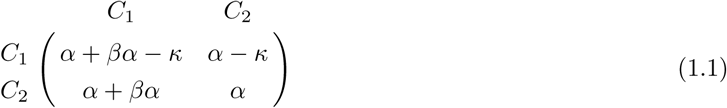

We assume that the linear benefit of the public good arises through growth factor diffusion that occurs on a timescale much faster than the average times between cell divisions. In a well-mixed population with fraction *u* of *C*_1_ cells, the fitness functions of this simple game are then be given by

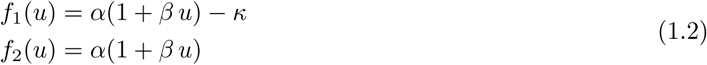

Our analysis in this manuscript is be based on cell type specific doubling rates, and in the case of logistic growth, also on the apoptotic rates. We are interested in the question when replicator dynamics, that typically only models changes in relative abundance as a result of fitness differences *f*_1_(*u*) − *f*_2_(*u*), predicts similar *ε*-fixation times as a logistic growth dynamics, and when this is not the case. The main idea is that the replicator dynamics neglects apoptotic rates, but that these rates in turn influence the time to reach an equilibrium in a co-growing and co-evolving heterogeneous cell population.

## 2 Methods

In this section we introduce our model of bounded frequency-dependent growth. We define our basic deterministic framework of two co-growing cancer cell populations, derive dynamic equations for the fraction of one clone and the total size of the population, and then derive an expression for the stable manifold of the system.

### 2.1 Logistic Growth Model

The population is assumed to consist of two types, and we denote their absolute numbers by *x*_1_ and *x*_2_. The carrying capacity is denoted by *K*, which we consider to be a constant. It possible to model it as a function of the strategies present in the population [28, 29]. The growth rate of each type is assumed to depend on the fraction of type 1 cells *u* = *x*_1_/(*x*_1_ + *x*_2_) according to growth functions *f*_1_(*u*) for type 1 and *f*_2_(*u*) for type 2. Lastly, cells of both types die at a constant rate *μ*. Taken together this implies that we get the following system of coupled logistic equations that describe co-growth and co-evolution of the two cell types:

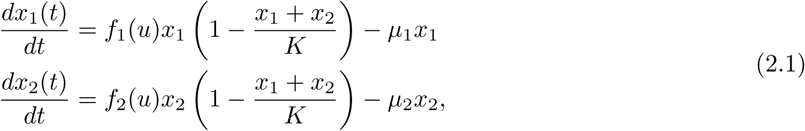

defined for *x*_1_*,x*_2_ ∈ ℝ^+^. In the following analysis we first assume *μ*_1,2_ = *μ* and *f*_1,2_(*u*) *> μ* for *u* ∈ [0,1], i.e. the net growth rate of both cells types will always be positive. In the second part of the discussion we will relax the assumption of equal rates and turn to the more general case of *α*_1_ ≠ *α*_2_, *μ*_1_ ≠ *μ*_2_, as we analyze the system implementing previously measured cellular rates of proliferation and apoptosis. Note that the logistic growth model emerges from a spatial setting that includes cell movement if cell migration occurs on a much faster time scale compared to cell division. It has been shown that in this case spatial correlations are negligible and the population dynamics can be described using a logistic growth equation [30]. In this parameter regime it is also justified to assume that interactions that influence the rate of cell division become independent of specific local configurations, and depend solely on the frequency of different cell types.

### 2.2 Analysis

In order to simplify the analysis of the system (2.1) we apply the following change of variables

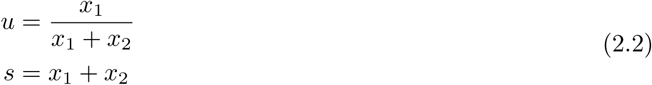

where *u* is the fraction of type 1 cells and *s* is the total population size. By differentiating *u* and *s* with respect to time we obtain the following system of ODEs

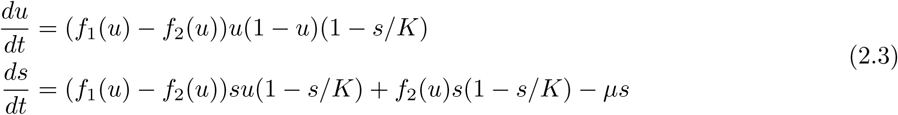

defined on *u* ∈ [0,1] and *s* ∈ ℝ^+^. We note that in the case when *s* is small compared to the carrying capacity *K*, such that *s/K* ≈ 0 the system reduces to

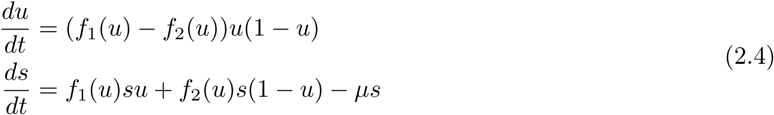

and we see that the equation for *u* is independent of the population size *s* and that *u* changes according to the standard replicator equation [13, 14]. We will now proceed to a more general analysis of our model.

#### Fixed points

By solving the equations

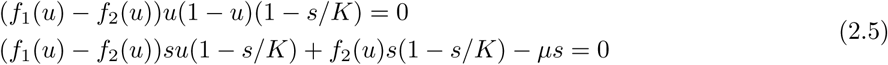

we see that for all growth functions *f*_1_ and *f*_2_ the system has the following set of fixed points on the boundary (see Appendix A for details):

1. (*u*_1_*,s*_1_) = (0,0) with corresponding eigenvalues *λ*_1_ = *f*_1_(0) − *f*_2_(0) and *λ*_2_ = *f*_2_(0) − *μ >* 0, which is unconditionally unstable
2. (*u*_2_*,s*_2_) = (1,0) with corresponding eigenvalues *λ*_1_ = *f*_1_(1) − *f*_2_(1) and *λ*_2_ = *f*_1_(1) − *μ >* 0, which is unconditionally unstable
3. (*u*_3_*,s*_3_) = (0*,K*(1 − *μ/f*_2_(0)) with corresponding eigenvalues 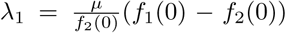 and *λ*_2_ = *μ* − *f*_2_(0) *<* 0, which is stable iff *f*_1_(0) *< f*_2_(0)
4. (*u*_4_*,s*_4_) = (1*,K*(1 − *μ/f*_1_(1)) with corresponding eigenvalues 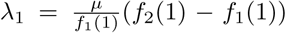 and *λ*_2_ = *μ* − *f*_1_(1) *<* 0, which is stable iff *f*_2_(1) *< f*_1_(1)

Here fixed point 1 and 2 are trivial in the sense that they correspond to a system void of cells. Fixed point 3 and 4 correspond to monoclonal populations and are stable if the resident type has a larger growth rate compared to the invading type.

If there are points *u*\* ∈ (0,1) such that *f*_1_(*u*\*) = *f*_2_(*u*\*), then these give rise to fixed points (*u*\**,K*(1 − *μ/*(*f*_1_(*u*\*)*u*\* + *f*_2_(*u*\*)(1 − *u*\*)))) which are stable if 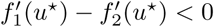 (see Appendix A for proof).

We note that the stability criteria for the non-trivial fixed points at *u* = 0 and 1, including potential internal fixed points, are identical with those of the two-type replicator equation with payoff functions *f*_1_ and *f*_2_.

#### Invariant manifold

We now focus our attention to the dynamics when the system is close to saturation (*s* ≈ *K*) with the aim of obtaining a simpler description of how the frequency *u*(*t*) changes in time. This can be achieved since the phase space contains a stable invariant manifold that connects all the non-trivial steady states. The invariant manifold is simply a curve *s* = *h*(*u*), which attracts the dynamics and once the system enters the manifold it will not leave it. This implies that the dynamics along the manifold are effectively one-dimensional, and can be captured with a single ODE for *u*(*t*).

If we write the invariant manifold as a function *s* = *h*(*u*), then, since it is invariant it must be tangent to the vector field 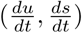 at every point. This implies the condition

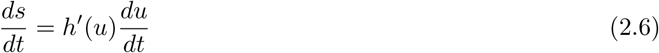

which is known as the manifold equation [31, 14]. By substituting 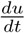 and 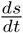 from (2.3) and letting *s* = *h*(*u*) we obtain the following equation for *h*(*u*)

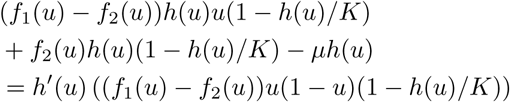

This equation is a non-linear ordinary differential equation and in order to solve it we express *h*(*u*) as a series expansion in the death rate *μ*, which typically is a small parameter

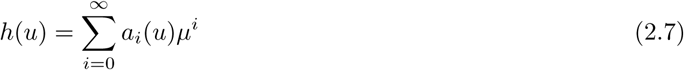

where *a*_*i*_(*u*) are coefficients that depend on *u*. We insert this ansatz into Eq. (2.6) and equate powers of *μ* to solve for the *a*_*i*_’s. We do this for *i* = 0,1,2, introduce 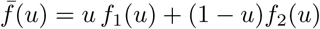, and get

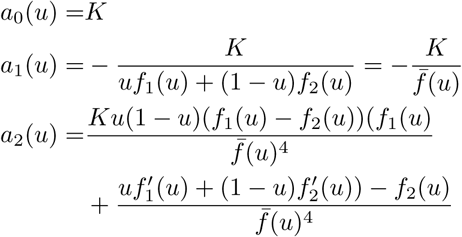

Numerical comparison shows that the invariant manifold is closely approximated by the first two terms, and we therefore drop all higher order terms and approximate the invariant manifold with

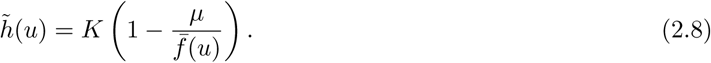

Note here that the complete solution would be more complicated, as can be seen from the fact that this first order expression does not solve the original manifold equation.

The dynamics along the invariant manifold are given by replacing *s* with 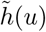 in (2.3), and we get the following expression (to first order in *μ*):

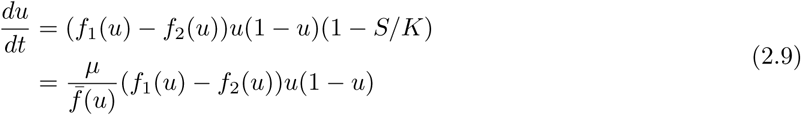

With the unusual pre-factor that is inversely proportional to the total fitness of the population, 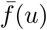, this equation for the frequency of type 1 cells is similar to the version of the replicator equation introduced my Maynard-Smith [32], and the one derived by Traulsen *et al.* [33] (if we disregard the demographic noise term). The difference compared to previous derivations is the factor *μ*, which implies that the rate of change of *u* along the invariant manifold is proportional to the death rate.

## 3 Results and Discussion

It is often argued that pre-factors to the replicator equation are irrelevant since the dynamic flow and fixed points remain unchanged. However, the time-scale of selection leading to an equilibrium might be altered. In this section we explore the difference between the standard replicator equation and the logistic model considered here. We examine this relationship in the context of a tumor-public goods game, in which some cells produce a public good at a cost, rendering a benefit to *all* cells in the population.

### 3.1 Diffusing public goods game

Autocrine production of growth factors is a common feature of cancer cells, and has previously been modeled using evolutionary game theory [23, 34]. Let us now consider two cell types that only differ in one aspect. Type 1 cells produce growth factor at a cost *κ*. Type 2 cells do not produce the growth factor and are termed free-riders. Otherwise, both cell types have the same growth rates, which are a linear function of growth factor availability. We assume that the growth factor production rate is given by *ρ* and that the growth factor is bound and internalized by both cell types at rate *δ*.

Two largely simplifying assumption are that, first, we are describing a well-mixed system and that, second, the growth factor concentration *G* is assumed to be uniform in space. We rely on the first assumption for mathematical convenience, as otherwise we would have to resort to non-analytical, agent based or hybrid modeling [35]. Secondly, additional growth factor provision was shown to be rapid and leading to high levels of tumor public good, provided that the respective genetic promoter was strong [27]. In a similar study by Cleary *et al.* [36], who studied Wnt1-based cooperative tumor evolutionary dynamics, aberrant expression of the cooperative signalling molecule was observed on a tumor wide scale. Thus, under these simplifying but productive assumptions, the growth factor dynamics obeys the equation

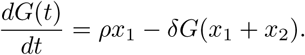

Further, we assume that the growth factor dynamics occur on a fast time scale compared to changes in *x*_1_ and *x*_2_. This implies that

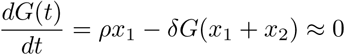

and we can solve for *G* to give

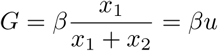

where *β* = *ρ/δ*. For simplicity, we first we consider a linear effect of the growth factor on the rate of cell division, as well as equal proliferation and death rates, which results in the growth functions given by Eqs. (1.2). In order for the growth rate to be larger than the death rate for all *u* we assume the inequality *α*−*κ > μ*. This choice of growth functions gives the following system of ODEs for the frequency of producers *u* and the total population size *s*:

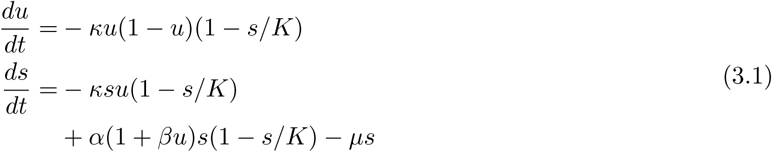

This system results from Eqs. (2.3) and has two non-trivial steady states given by a monomorphic population of free-riders (0,1−*μ/α*), and a population consisting only of producers (1,1−*μ/*(*α*(1+*β*)−*κ*)), see analysis following Eqs. (2.5). The eigenvalues are

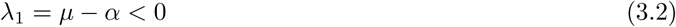

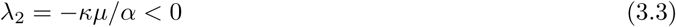

and hence the free-rider steady state is stable. For the other fixed point (producers dominate) we have

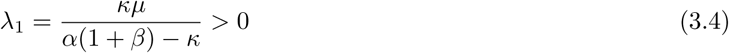

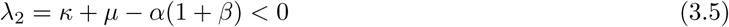

making it unstable. Figure 1 A shows the phase space of the system, where the open circles indicate unstable steady-states and the filled circle shows the location of the single stable steady state. We note that for almost all initial conditions the dynamics rapidly converge to the invariant manifold (2.8) which is approximately given by

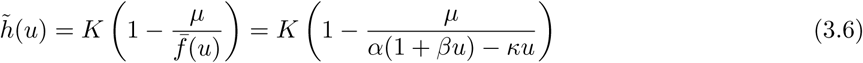

Once the system enters the invariant manifold the dynamics can be approximated by (2.9) which for the diffusing public goods game considered here are given by

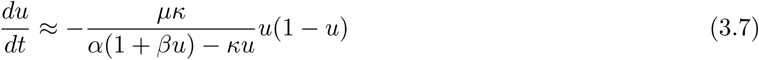

Thus, in order to assess the impact of cell death and turnover on selection, we compare our description of the public goods game (3.1) with the standard replicator equation

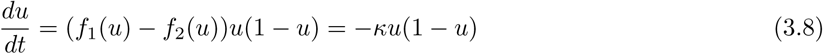

**Figure 1.**
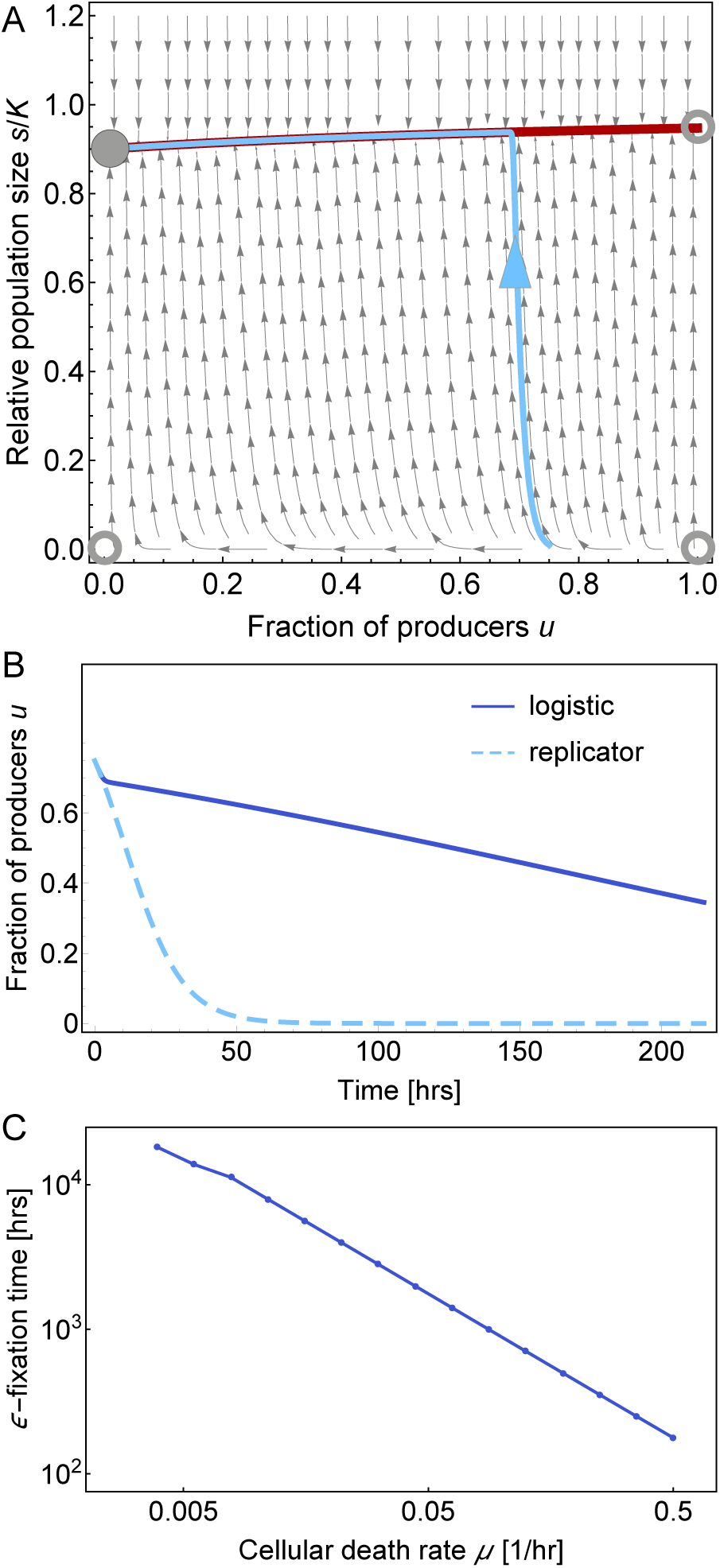
(A) Phase space of the ODE-system (3.1) describing the diffusing public goods game. The gray arrows show the flow lines of the system, the open circles show the three unstable stationary states, and the filled circle shows the only stable steady states where the population is dominated by non-producing type 2 cells. The red line shows the invariant manifold (3.6), and the light blue curve (with arrow pointing forward in time) shows one solution of the deterministic system as it approaches and eventually follows the stable manifold. (B) The frequency of producers *u*(*t*) obtained from the logistic system and the standard replicator equation (the line is just a guide to the eye). (C) The *ε*-fixation time measured as the time it takes to reach the state *u* = 0.001. In all panels, the values are *α* = 1.0, *β* = 1.0, *κ* = 0.1, *K* = 1, and *μ* = 0.1 (A,B), where we chose to observe time in units of hours. The initial conditions are (*u*_0_*,s*_0_) = (0.75,0.01 ∗ *K*).

Figure 1 B shows a comparison between the solution of the logistic system (3.1) and the replicator equation (3.8) for the same initial condition *u*_0_ = 0.75 (*s*_0_ = 0.01*K*) and with a death rate of *μ* = 0.1/hour. Whereas the two solutions agree for small times (when *s* ≪ *K*), they start to diverge as soon as the solution to the logistic system enters the invariant manifold. The solution of the replicator equation quickly converges to the steady state *u* = 0, while the fraction of producers in the logistic case decreases approximately linearly with time.

In order to quantify the effect of the death rate *μ* on the rate of selection we measured the time it takes for the logistic system to approach a steady state. For a fixed intial condition (*u*_0_*,s*_0_) = (0.75,0.01) we measured the time it took for the system to reach a small *ε* neighbourhood of the fixed point, i.e. |*u*(*t*)−*u*\*|≤ *ε*, with *u*\* = 0 and *ε* = 0.01. We call this time the *ε*-fixation time. All other parameters were fixed at *α* = *β* = 1, *κ* = 0.1, *μ* = 0.1 and *K* = 1. The result is displayed in Figure 1 C and shows that the *ε*-fixation time scales as *μ*^−1^. This implies that for small *μ* the time it takes the system to reach the steady state can be exceedingly long. It is worth noting that the *ε*-fixation time for the replicator equation can be obtained in the limit of 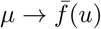, performed on the logistic system, implying a never-growing population, in which the death rate equals the average birth rate.

### 3.2 Timescales of *in vivo* and *in vitro* cellular expansions

Previous studies of ecological interactions in growing tumor cell populations have observed various forms of frequency-dependent effects. These effects have then been linked to the persistence of distinct cancer cell lines that provide growth enhancing public goods to the tumor, most notably in experimental work by Marusyk *et al.* [27]. There, it could be shown that a mixture of certain clones could not explain tumor outgrowth in vivo by simply using superposition of individual clonal birth and death rates. Rather, synergistic tumor-driving effects can emerge, pointing to more intricate, potentially frequency-dependent growth effects, based on direct or indirect clonal interactions [18]. For the purpose of illustration, we extracted individual clonal proliferation (*α*_*i*_) and death rates (*μ*_*i*_) from Marusyk *et al.* [27], in order to predict how these rates shape the dynamics. Out of 16 clonal cell lines, each distinctively expressing a different gene, we chose four clones to calculate baseline cellular birth and death rates. The four clones, derived from the breast cancer cell line MDA-MB-468, were LoxL3 (lysyl oxidase type 3 [37], linked to breast cancer invasion and metastasis), IL11 (interleukin 11, a member of the IL 6 family that plays a multifaceted role in leukemia and breast cancer [38]), and CCL5 (C-C motif ligand 5, a chemokine with emerging roles in immuno-therapy [39]). The baseline cellular birth and death rates of these clones were calculated in the following way, based on *in vivo* growth experiments, originally performed in a mouse xenograft model (“tumors formed by orthotopic trans-plantation into the mammary fat pads of immunodeficient Foxn1^nu^ (nu) mice” [27]). For all four clones, it was established that tumors grew exponentially; from longitudinal measurements and associated cellularity calculations, the net cellular doubling rates were calculated (see Ext. Data Fig. 3 and SI in [27], where exponential growth rates are given, which we transformed into doubling rates). For the four above mentioned clones, proliferation assays were also performed (Ext. Data Fig. 1 in [27]). These BrdU staining experiments measure the fraction of cells in S-phase of the cell cycle, *χ*. S-phase duration *T*_S_ is highly conserved in mammary cells [40], known to be about 8 hours long, *χ* serves as a direct estimate for the percent of S-phase in relation to the whole cell cycle *T*, and thus the doubling rate, which we set to *α* = 1*/T*. Using the relation

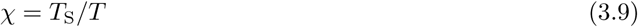

we calculated the mono-clonal birth rates using

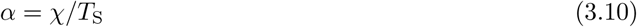

Thus, given the net doubling rate *r* = *α* − *μ*, it is possible to estimate the death rate

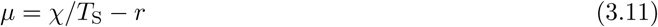

with *T*_S_ fixed to 8 hours. Data for *r* and *χ* are given in Appendix B. Since for both *r* and *χ*, several independent measurements were performed, we calculated distributions of *α* and *μ* for the three cell lines described above. We contrasted these distributions to *in vitro* distributions of cellular birth and death rates, adapted from [41] (Fig. 3 therein), which are, notably, very similar to other *in vitro*-values, e.g. reported for the PC-9 non-small cell lung cancer cell line [42], see Figure 2 A. In the *in vivo* tumor growth experiments, exponential growth was observed within the time frame of 50 to 80 days, at growth rates up to two population doublings per day (net growth rate) [27]. However, in most tumors the net growth rate was more moderate, and the actual cellular birth and death rates were at least of similar order in magnitude (*α/μ* ≈ 1). This stands in contrast to the birth-death rate ratios observed in cell cultures, where birth rates often exceed death rates by an order of magnitude (*α/μ* ≈ 10) [42, 43, 41, 44].

**Figure 2.**
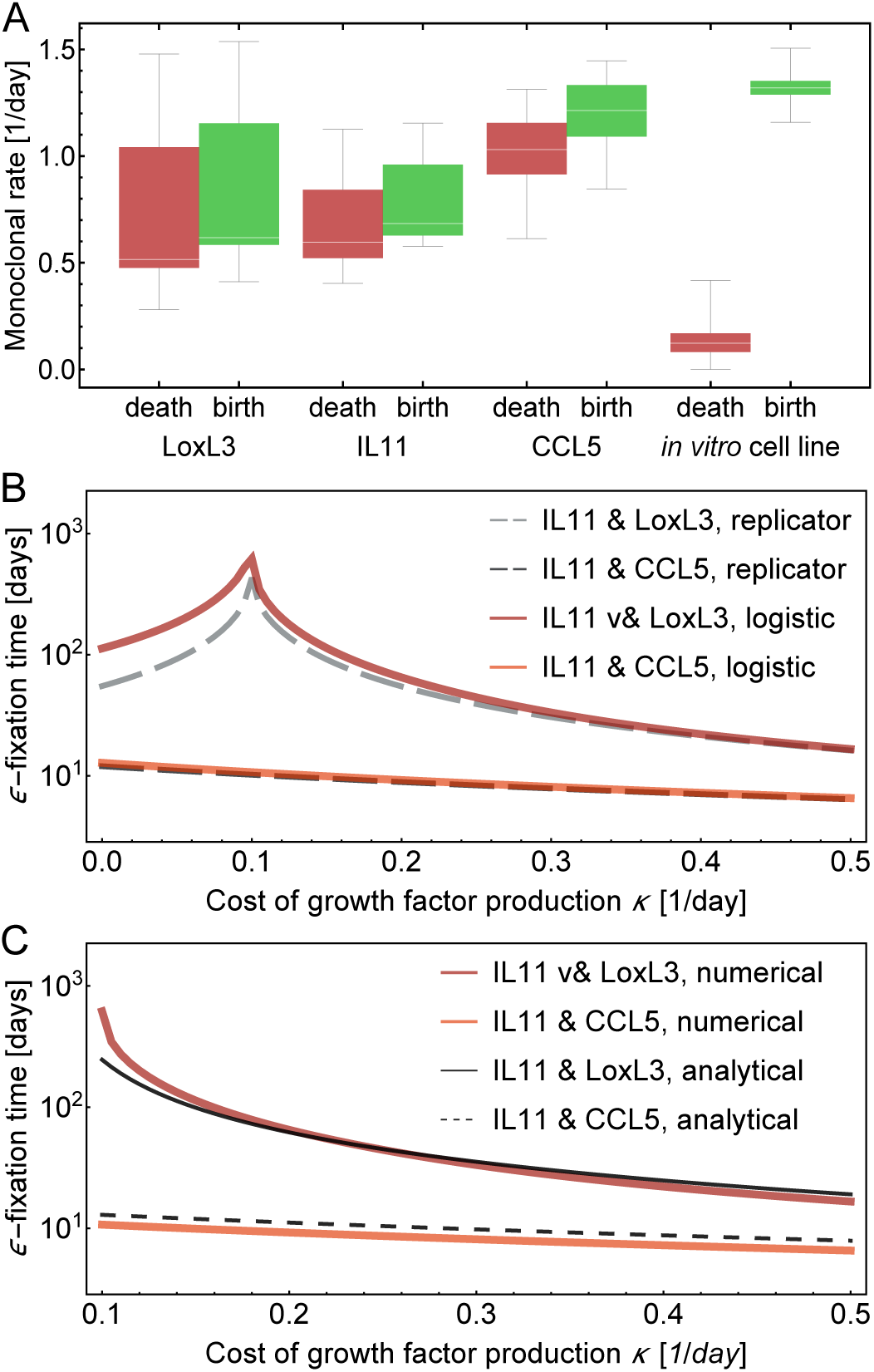
(A) Birth and death rate distributions, calculated from previous experiments, where engineered breast cancer cell lines, characterized over-expressing certain cytokines, were observed to grow in *in vivo* xenograft mouse model tumors [27]. Although net tumor growth was high, death and birth rates were similar in all clones considered. In comparison, we also show *in vitro* cell line rates, estimated by Juarez et al. [41]. We further used the fact that the IL11 cells are growth factor producers. (B) Using median birth and death rates from the distributions in (A), we measured the *ε*-fixation time numerically determined using Eqs. (3.13) (defined as the time to reach an *ε*-neighborhood equilibrium value of *u*, with *ε* = 0.001, *u*_0_ = 0.5) and compared it to the *ε*-fixation time of the standard replicator equation (3.8). Note that we used Eqs. (3.13) for this numerical procedure. For IL11 we used *α*_1_ = 0.684/day and *μ*_1_ = 0.596/day. For LoxL3 we used *α*_1_ = 0.617/day and *μ*_1_ = 0.515 /day For CCL5 we used *α*_1_ = 1.214/day and *μ*_1_ = 1.031/day. *β* = 1, with *u*_0_ = 0.5 and *s*_0_ = 0.01*/K*. Note here that the peak in *ε*-fixation time marks the shift from *u* → 1 to *u* → 0 as the cost increases; this transition can only occur when producers and non-producers have similar birth and death rates. (C) Comparison of *ε*-fixation times determined numerically using (3.13) to the analytical approximation (3.19), parameters the same as in (B).

As a notable difference to the previous chapter, here we assume both *α*_1_ ≠ *α*_2_ and *μ*_1_ ≠ *μ*_2_. Thus, instead of (2.3), we now deal with the more general payoff structure

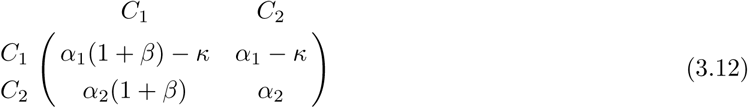

and obtain the following ODEs for frequency of producer cells and total size of the system

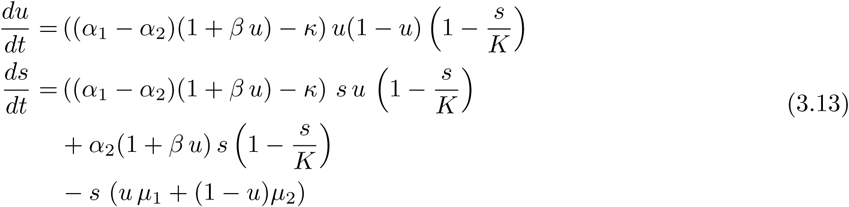

and we seek to estimate the time it takes to reach a small *ε* neighborhood of the equilibrium |*u*(*t*)−*u*\*|≤ *ε*, shown in Figure 2 B. The combinations IL11 with one other cell line was chosen because it has been established that IL11 is a growth factor producer clone, which, at least in a first approximation, renders a linear fitness benefit [27]. We here make the additional assumption that IL11 cells carry a cost associated with growth factor production, and explore the extinction process of IL11 cells as they compete with either CCL5 or LoxL3 cells (Figure 2).

We can calculate an estimate of the ‘time to fixation’ in the following way. Suppose the fraction of growth factor producers, *u*, is at a stable equilibrium, and that there are only two possible stable equilibria, *u*\* = 0 and *u*\* = 1. Then, the stationary solutions for the population size, *s*\*(*u*\*), will be

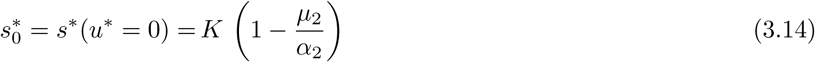

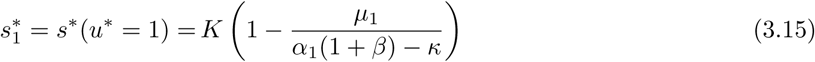

We now assume that the total population size remains at the stationary value, although it in fact changes (slightly) with *u*. This assumption can be thought of as a zeroth-order approximation in 1 − *s/K*, and it implies that near the stable manifold, the frequency *u* obeys the ODE

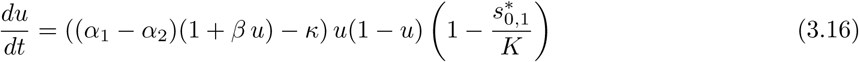

which we can solve by inserting the approximations (3.14) and (3.15) into the ODE (3.16) and get the two solutions (for two different possible endpoints)

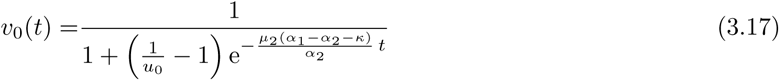

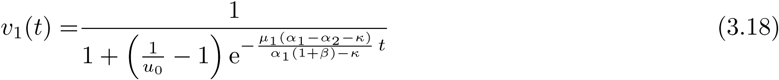

We now seek solutions of 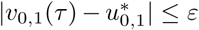 for *τ* (with the equilibrium points 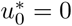, 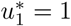), and find the following relations that approximate the *ε*-fixation times

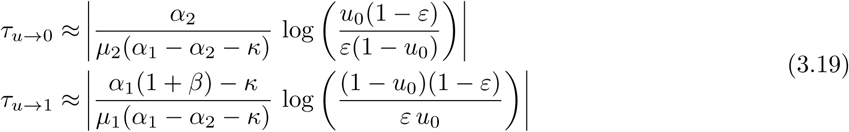

where *u*_0_ is the initial frequency. Note that here, we deviate from the notion of (average) *ε*-fixation times in the strict stochastic sense [45], and replace the term by a threshold-based analytical approximation. Especially in a population that has reached the stable manifold, even a small fraction of remaining producers cells could still mean that there are as many cells as needed to warrant a mean-field rather than a fully stochastic description.

For the *u* → 0, *s* → *K*(1 − *μ*_2_*/α*_2_) case, we can now compare our analytical approximations with the *ε*-fixation times of the full numerical solution in Figure 2 C, as a function of *κ*. Depending on the differences in clonal birth and death rates, the approximation exhibits qualitative differences. Eq. (3.19) consistently overestimates the *ε*-fixation time if the death rate of the producer cells is lower than that of non-producers (IL11 with CCL5 *α*_1_ − *μ*_1_ *< α*_2_ − *μ*_2_), but it underestimates the *ε*-fixation time if the net growth rate of the producer cells is higher than that of non-producers as long as the cost of growth factor production does not exceed a certain threshold (IL11 with LoxL3, *α*_1_ − *μ*_1_ ≈ *α*_2_ − *μ*_2_). Hence, not only the cost of growth factor production factor influences the time to extinction of producer cells, also the monoclonal net growth rate influences both the time to extinction of producers and the impact of an assumed cost associated with growth factor production. The approximations (3.19) are of ‘zero-order’ in changes in *s*. Yet, they are still able to reflect the basic fact that *ε*-fixation time can be heavily influenced by the cellular death rate of the abundant cell type. According to our rough approximation, the extinction time of producer cells (3.19) is both proportional to the ratio of birth to death rate of the non-producers, as well as inversely proportional to the birth rate difference. Surprisingly, in this approximation *τ*_*u*→0_ does not depend on the absorption or production rate of the growth factor, captured by *β*. Large differences in baseline birth rates extend growth-factor producer extinction times. For larger values of *α*_2_*/μ*_2_, the extinction time is less sensitive to changes in the cost of growth factor production.

The two cellular death rates *μ*_1_ and *μ*_2_ have different effects on *ε*-fixation times. We used numerical solutions of the full system (3.13), in comparison to the replicator equation (3.8), to analyze variability of *ε*fixation times (extinction of growth factor producer cells) under variable individual death rates. Thereby, we recover that higher total death rate speeds up the *ε*-fixation time across different initial conditions (Figure 3 A), and that the death rate of the ‘winner-clone’ plays a more important role (Figure 3 B): *μ*_2_ has a more pronounced impact on the *ε*-fixation time of non-producers. This might be connected to the fact that apoptosis-driven cell turnover of the nearly dominant cell type (i.e. the non-producer cells) governs the *ε*-fixation time. In accordance with this observation, the stable manifold itself governed by the apoptotic rate of the dominant clone, compare to Eq.(2.8).

**Figure 3.**
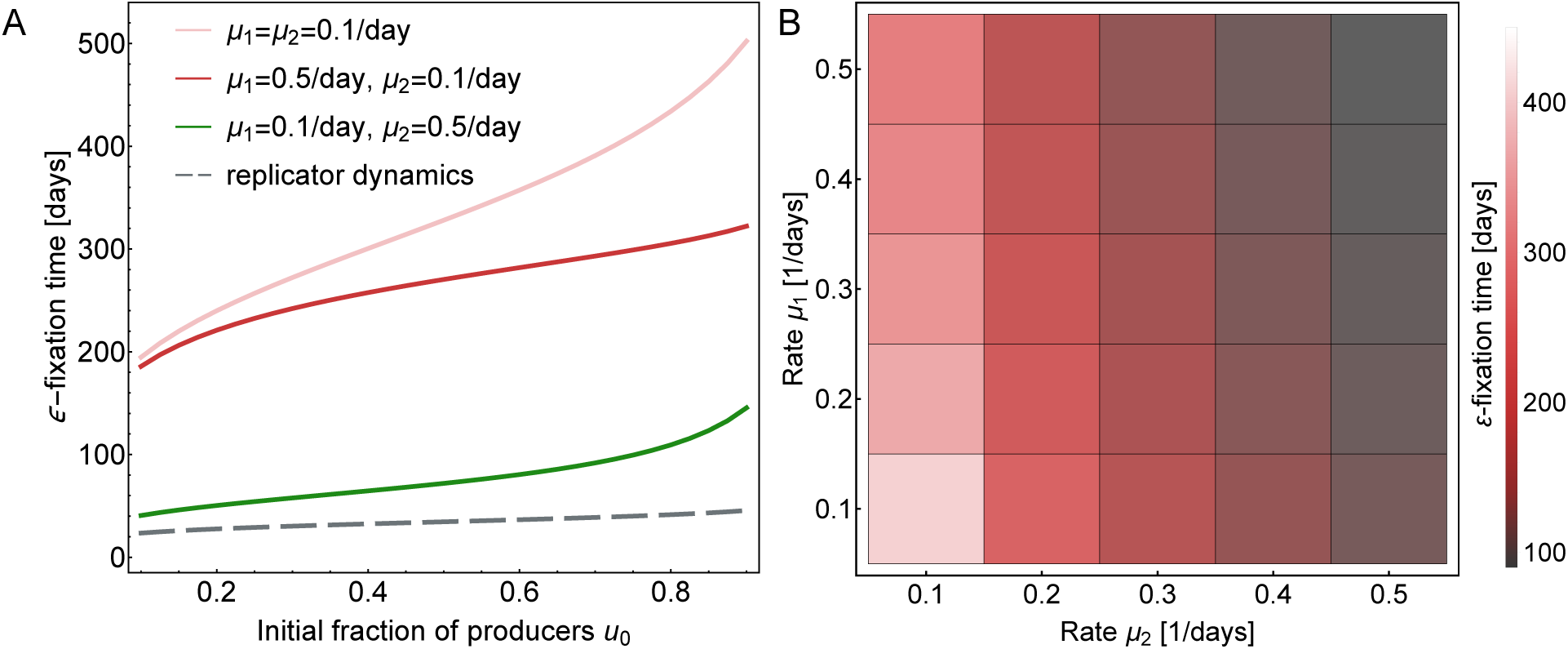
(A) Comparing the influence of the death rate of producers, *μ*_1_ with the influence of the death rate of non-producers, *μ*_2_. (B) Variation of the extinction time of growth factor producers under different death rates *μ*_1_ and *μ*_2_, *u*_0_ = 0.8. In all panels, *κ* = 0.2, *β* = 1, *α*_1,2_ = 1.0/day, *s*_0_ = 0.01 ∗ *K*, and *K* = 1. The times here were calculated by numerical integration of Eqs. (3.13). An observation that cannot be explained with our analytical approximations is that the extinction times of the logistic growth dynamics tend to be closer to the (short) extinction times of the replicator dynamics for smaller death rate *μ*_1_ (*μ*_2_ fixed). However, the utility of our approximations is supported by the observation that overall variability in *ε*- fixation times is driven by *μ*_2_, the death rate of the dominant cell type.

## 4 Summary and Conclusions

We here have presented calculations that were concerned with the stability and time to reach a neighborhood of equilibrium points in evolutionary game dynamics between two types of tumor cells. We focused on the dynamics of a tumor public good (tumor growth factor), in which we assumed linear fitness functions of growth factor producers and non-producers. The fitness function linearly depends on the relative abundance of growth factor producers, and production comes at a cost. We did not need assume that the evolving population was at carrying capacity, as reflected in the logistic growth model. Thus, in general, population expansion and cellular birth as well as death rates are of importance for the time the system takes to equilibrate. The standard replicator equation typically rules out explicit death effects, and thus may not accommodate the impact of these death rates on the time to reach a population equilibrium.

The use of replicator equations and birth-death processes assume constant population size [7] or a population which is growing uniformly, for example at an exponential rate [13]. These assumptions have led to a plethora of fruitful results in evolutionary game theory [46], e.g. to the ability to understand fixation and extinction times in evolutionary 2×2-games [47, 48, 49, 50], multiplayer-games [51], structured populations [52], or bi-stable allelic competition [53, 54]. Evolutionary games have also been used to establish rules for equilibrium selection even in complex group-coordination games [55, 56], in chemical game theory [57], and to map complex tumor dynamics [58, 59, 60, 25, 61, 23, 34, 24]. However, the assumption that the population is either at constant size may be limiting, as also recently discussed by Li *et al.* in the context of co-growing and co-evolving bacterial species [62]. Instead, the near-equilibrium population size and the time to reach equilibria are influenced directly by birth and death rates in the population.

We show that, for small differences between the birth and death rates, the eco-evolutionary dynamics of the mixture of two clones may be approximated by standard replicator dynamics. Analysis of previously established growth-factor dependent tumor dynamics of *in vivo* tumor growth showed that this parameter regime might indeed be biologically relevant (Figure 2), even when the tumor population has not reached its carrying capacity. However, prominent examples of *in vitro* cell line expansions demonstrate that large differences between cellular death and birth rates might impact the dynamics in a different way [42, 43, 44], and in this case the replicator equation is a poor approximation of the eco-evolutionary dynamics. We used a logistic growth model that includes cell death. This system describe both co-growth, as well as co-evolution of two tumor cell types. The choice of logistic growth is by no means unique, but a simple, first-order form of non-uniform growth.

We report two major findings. First, a first order approximation in death rates allows estimation of the stable manifold, and reveals linear dependence on the apoptotic rate of the more abundant cell type. Second this knowledge can be used to inform a zero-order approximation (in constant system size) of the time to get arbitrarily close to equilibrium (‘*ε*-fixation time’), which reveals that indeed the cellular turnover of the dominant cell type near equilibrium governs the*ε*-fixation time as the system slowly moves along stable manifold. This framework allowed us to examined the degree of the resulting variability in *ε*-fixation times based on previously measured *in vivo* tumor cell proliferation and death rates in the context of competition between producers and non-producers of a growth-factor public good.

Various aspects of cancer cell population structure, such as cellular differentiation, localization or spatial heterogeneity, point to dynamic non-linear size changes over time, especially during treatment [63, 64, 65, 66], and treatment can shift the evolutionary game [67]. Furthermore, selection mechanism that go beyond relative fitness differences play a role in our understanding of other biological and clinically relevant systems, such as the hematopoietic system [68, 69]. Hence, future modeling efforts that seek to apply evolutionary game theory to explain complex cancer growth patterns need to precisely disentangle complex interaction patterns between cells from the overall growth kinetics of a tumor. Detailed understanding of tumor growth kinetics is especially important in co-growing populations, as we here show that the convergence towards an equilibrium–which sets the time scale for potential treatment and relapse effects–sensitively depends on the microscopic cellular growth rates. The often performed, and mathematically convenient re-scaling of time that leads to replicator equations might eliminate effects that are crucial for understanding transitions between equilibria and describing relevant time scales of tumor evolution.

## Author contributions

All authors conceived the study, analyzed the data, performed mathematical and statistical modeling, and wrote the paper.

## Acknowledgements

We thank Joel S. Brown (Chicago) and Christian Hilbe (Vienna) for valuable comments and suggestions.

## Data Accessibility

All data used in this manuscript are presented in the Appendix and can be found online with the cited references.

## A Fixed points and stability

In order to investigate the stability of the fixed points of (2.3) we denote the right hand sides by:

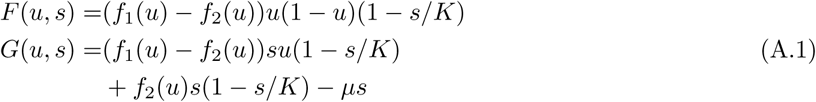

and calculate the Jacobian at the fixed point (*u*\**,s*\*)

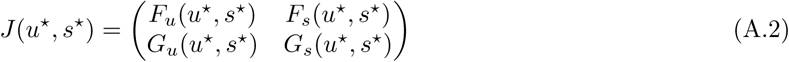

where subscript denotes partial derivative with respect to *u* and *s*.

### Boundary fixed points

For the boundary fixed points we find the following:

At (*u*\**,s*\*) = (0,0) we find that

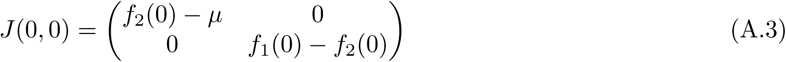

with eigenvalues *λ*_1_ = *f*_1_(0) − *f*_2_(0) and *λ*_2_ = *f*_2_(0) − *μ >* 0. The last inequality holds because we assumed a positive net growth rate for both cell types for all *u* ∈ [0,1]. This fixed point is therefore unconditionally unstable.

At (*u*\**,s*\*) = (0,1) we find that

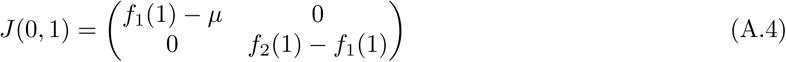

with eigenvalues *λ*_1_ = *f*_1_(1)−*f*_2_(1) and *λ*_2_ = *f*_1_(1)−*μ >* 0. Again, the inequality holds because we assumed a positive net growth rate for both cell types for all *u* ∈ [0,1]. This fixed point is therefore unconditionally unstable.

At (*u*\**,s*\*) = (1*,K*(1 − *μ/f*_1_(1)) we find that

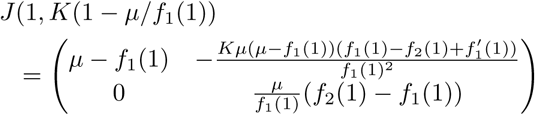

with eigenvalues 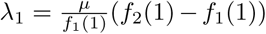 and *λ*_2_ = *μ*−*f*_1_(1) *<* 0. This implies that the fixed point is stable iff *f*_2_(1) *< f*_1_(1).

At (*u*\**,s*\*) = (0*,K*(1 − *μ/f*_2_(0)) we find that

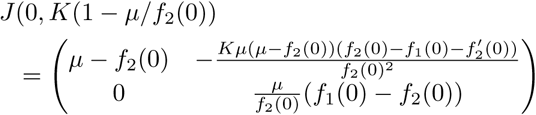

with eigenvalues 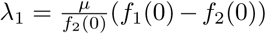 and *λ*_2_ = *μ*−*f*_2_(0) *<* 0. This implies that the fixed point is stable iff *f*_1_(0) *< f*_2_(0).

### Internal fixed points

Internal fixed points exist at points where *f*_1_(*u*\*) = *f*_2_(*u*\*) for 0 *< u*\* *<* 1. The corresponding *s*- coordinate is given by solving 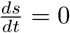 in terms of *u* to get *s*\* = *K*(1 − *μ/f*_1_(*u*\*)). The Jacobian at such a point is given by

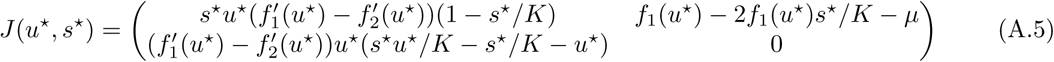

In order to say something about the stability of such a point we need to investigate the signs of the eigenvalues of *J*. We do this by looking at the sign of each matrix entry. For now, we assume nothing about the sign of 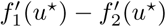 and instead focus on the other factors in each matrix entry.

First we see that

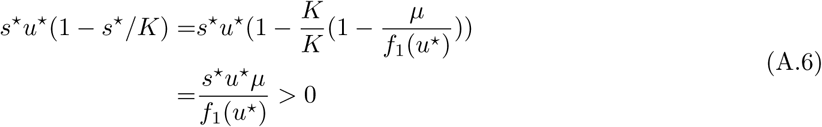

Further we have

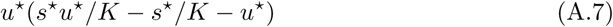

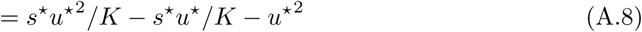

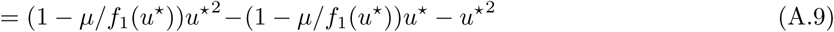

Here 0 ≤ (1 − µ*/f*_1_(*u\**)) *<* 1 since *f*_1_ (*u*) > *µ ≥* 0. This implies that

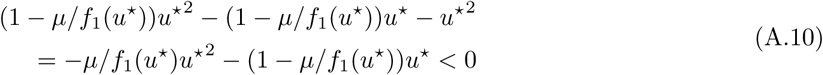

since both terms are negative. Lastly we see that

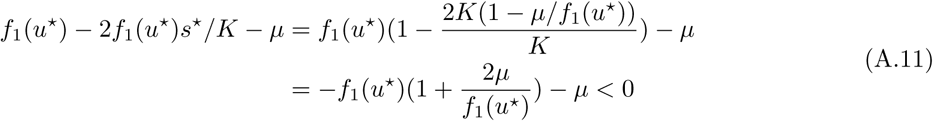

since 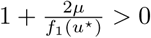

This implies that we can write the Jacobian as

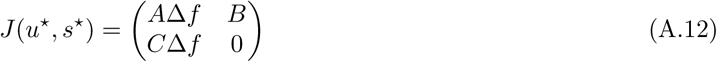

where *A >*0, *B <*0, *C <*0 and 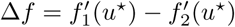. The eigenvalues of the Jacobian are given by

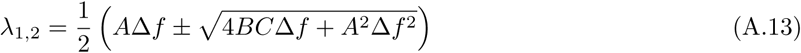

Now if Δ*f* > 0 then *A*Δ*f* > 0, and 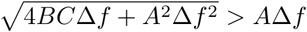. This implies that *λ*_1_ *>* 0 and *λ*_2_ *<* 0, and hence the fixed point (*u*\**,s*\*) is unstable.

If on the other hand Δ*f <* 0 then there are three possibilities, either (i) 4*BC*Δ*f* + *A*^2^Δ*f*^2^ *>* 0 or (ii) 4*BC*Δ*f* + *A*^2^Δ*f*^2^ *<* 0 or (iii) 4*BC*Δ*f* + *A*^2^Δ*f*^2^ = 0. If (i) holds then 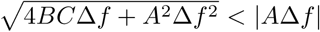 which implies that *λ*_1,2_ *<* 0. If (ii) is the case then 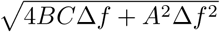 is complex and <(*λ*_1,2_) *<* 0. Lastly if (iii) is the case then *λ*_1,2_ = *A*Δ*f/*2 *<* 0.

This shows that the stability of the stationary point at (*u*\**,s*\*) is fully determined by the sign of 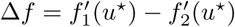. If Δ > 0 the point is unstable and if Δ*f <* then the point is stable.

## B Clonal population doubling rates

Here, all rates are given per day; *in vivo* data taken from Marusyk *et al.* [27].

For LoxL3, we used the following population doubling rates (net growth rates)

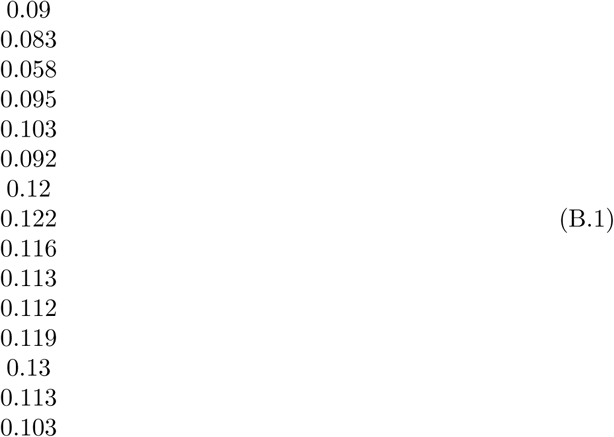

and the following percentage of S-phase during cell cycle *χ*

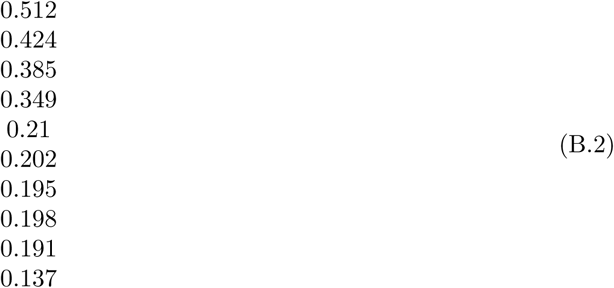

For IL11, we used the following population doubling rates (net growth rates)

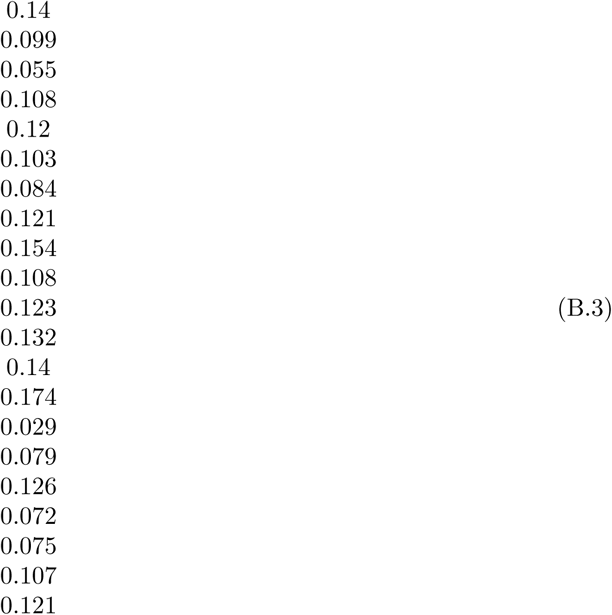

and the following percentage of S-phase during cell cycle *χ*

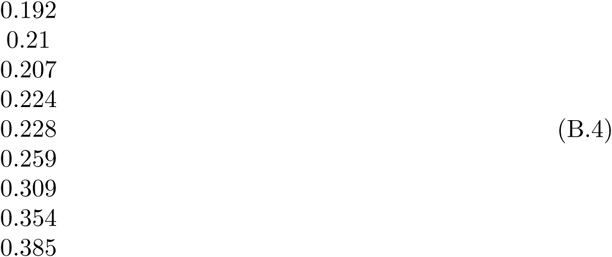

For CCL5, we used the following population doubling rates (net growth rates)

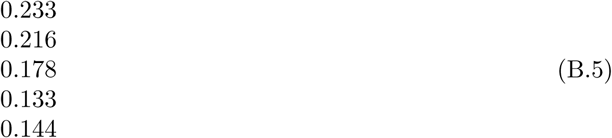

and the following percentage of S-phase during cell cycle *χ*

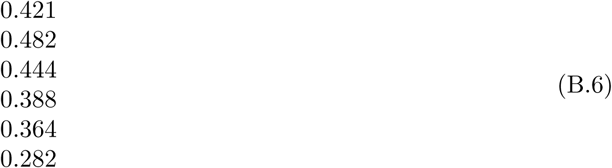

The distributions shown in Figure 2 resulted from all possible pairs of these numbers to calculate *α* and *μ*, Eqs. (3.10) and (3.11).

For generation of the *in vitro* distributions we used normally distributed rates (truncated by 0), with a mean death rate of 0.12/day (SD 0.0672) and a mean birth rate of 1.32/day (SD 0.048), adapted from Juarez et al. [41].

## References

1. von Neumann J, Morgenstern O. Theory of Games and Economic Behavior. Princeton: Princeton University Press; 1944.

2. Aumann RJ. Game Theory. In: Milgate EM, Newman P, editors. The New Palgrave: A Dictionary of Economics. vol. 2. London, UK: Macmillan; 1987. p. 460.

3. Ross D. Game Theory. In: Zalta EN, editor. The Stanford Encyclopedia of Philosophy (Fall 2010 Edition); 2010. p. URL: http://plato.stanford.edu/archives/fall2010/entries/game-theory.

4. Nash JF. Equilibrium points in n-person games. Proceedings of the National Academy of Sciences USA. 1950;36:48–49.

5. Myerson RB. Game Theory. Analysis of Conflict. Harvard University Press, Cambridge, MA; 1997.

6. Vincent TL, Brown JS. Evolutionary Game Theory, Natural Selection, and Darwinian Dynamics. Cambridge University Press, Cambridge UK; 2005.

7. Nowak MA. Evolutionary Dynamics. Cambridge MA: Harvard University Press; 2006.

8. Gintis H. Game Theory Evolving. Princeton: Princeton University Press; 2000.

9. Maynard Smith J, Price GR. The logic of animal Conflict. Nature. 1973;246:15–18.

10. Traulsen A, Pacheco JM, Nowak MA. Pairwise comparison and selection temperature in evolutionary game dynamics. Journal of Theoretical Biology. 2007;246:522–529.

11. Sandholm WH. Pairwise comparison dynamics and evolutionary foundations for Nash equilibrium. Games. 2010;1(1):3–17.

12. Wu B, Bauer B, Galla T, Traulsen A. Fitness-based models and pairwise comparison models of evolutionary games are typically different–even in unstructured populations. New Journal of Physics. 2015;17:023043.

13. Taylor PD, Jonker L. Evolutionarily stable strategies and game dynamics. Mathematical Biosciences. 1978;40:145–156.

14. Hofbauer J, Sigmund K. Evolutionary Games and Population Dynamics. Cambridge University Press, Cambridge; 1998.

15. Sandholm WH. Population games and evolutionary dynamics. MIT Press, Cambridge, MA; 2010.

16. Milinski M. Tit For Tat in sticklebacks and the evolution of cooperation. Nature. 1987;325(6103):433– 435.

17. Szorkovszky A. Evolutionary game theory: a case of too much theory? Blog: Collective Behaviour. 2015 October;Available from: http://www.collective-behavior.com/ evolutionary-game-theory-a-case-of-too-much-theory [cited 2017].

18. Tabassum DP, Polyak K. Tumorigenesis: It takes a village. Nature Reviews Cancer. 2015;15:473–483.

19. Junttila MR, de Sauvage FJ. Influence of tumour micro-environment heterogeneity on therapeutic response. Nature. 2013;501(7467):346–354.

20. Scott J, Marusyk A. Somatic clonal evolution: A selection-centric perspective. Biochimica et Biophysica Acta (BBA)-Reviews on Cancer. 2017;.

21. Marusyk A, Almendro V, Polyak K. Intra-tumour heterogeneity: a looking glass for cancer? Nature Reviews Cancer. 2012;12(5):323–334.

22. Altrock PM, Liu LL, Michor F. The mathematics of cancer: integrating quantitative models. Nature Reviews Cancer. 2015;15:730–745.

23. Archetti M, Ferraro DA, Christofori G. Heterogeneity for IGF-II production maintained by public goods dynamics in neuroendocrine pancreatic cancer. Proceedings of the National Academy of Sciences USA. 2015;112:1833–1838.

24. Kaznatcheev A, Vander Velde R, Scott JG, Basanta D. Cancer treatment scheduling and dynamic heterogeneity in social dilemmas of tumour acidity and vasculature. British Journal of Cancer. 2017;116:785–792.

25. Kianercy A, Veltri R, Pienta K. Critical transitions in a game theoretic model of tumour metabolism. Interface focus. 2014;.

26. Tomlinson I, Bodmer W. Modelling the consequences of interactions between tumour cells. British Journal of Cancer. 1997;75(2):157.

27. Marusyk A, Tabassum DP, Altrock PM, Almendro V, Michor F, Polyak K. Non-cell-autonomous driving of tumour growth supports sub-clonal heterogeneity. Nature. 2014;514:54–58.

28. Novak S, Chatterjee K, Nowak MA. Density games. Journal of Theoretical Biology. 2013;.

29. Gerlee P, Anderson AR. The evolution of carrying capacity in constrained and expanding tumour cell populations. Physical biology. 2015;12(5):056001.

30. Baker RE, Simpson MJ. Correcting mean-field approximations for birth-death-movement processes. Physical Review E. 2010;82:041905.

31. Wiggins S. Introduction to applied nonlinear dynamical systems and chaos. Springer Science & Business Media, New York, NY; 2003.

32. Maynard Smith J. Evolution and the Theory of Games. Cambridge University Press, Cambridge; 1982.

33. Traulsen A, Claussen JC, Hauert C. Coevolutionary dynamics in large, but finite populations. Physical Review E. 2006;74:011901.

34. Gerlee P, Altrock PM. Complexity and stability in growing cancer cell populations. Proceedings of the National Academy of Sciences USA. 2015;112:E2742–E2743.

35. Rejniak KA, Anderson ARA. Hybrid models of tumor growth. Reviews in System Biology and Medicine. 2011;3:115–125.

36. Cleary AS, Leonard TL, Gestl SA, Gunther EJ. Tumor cell heterogeneity maintained by cooperating subclones in Wnt-driven mammary cancers”. Nature. 2014;508:113–117.

37. Molnar J, Fong KS, He QP, Hayashi K, Kim Y, Fong SF, et al. Structural and functional diversity of lysyl oxidase and the LOX-like proteins. Biochimica at Biophysica Acta. 2003;1647:220–224.

38. Johnstone CN, Chand A, Putoczki TL, Ernst M. Emerging roles for IL-11 signaling in cancer development and progression: Focus on breast cancer. Cytokine & Growth Factor Reviews. 2015;26:489–498.

39. Halama N, Zoernig I, Berthel A, Kahlert C, Klupp F, Suarez-Carmona M, et al. Tumoral Immune Cell Exploitation in Colorectal Cancer Metastases Can Be Targeted Effectively by Anti-CCR5 Therapy in Cancer Patients. Cancer Cell. 2016;29:587–601.

40. Bell SP, Dutta A. DNA Replication in Eukariotic Cells. Annual Reviews in Biochemistry. 2002;71:333–374. 19

41. Juarez EF, Lau R, Friedman SH, Ghaffarizadeh A, Jonckheere E, Agus DB, et al. Quantifying differences in cell line population dynamics using CellPD. BMC Systems Biology. 2016;10:92.

42. Chmielecki J, Foo J, Oxnard GR, Hutchinson K, Ohashi K, Somwar R, et al. Optimization of Dosing for EGFR-Mutant Non-Small Cell Lung Cancer with Evolutionary Cancer Modeling. Science Translational Medicine. 2011;3:90ra59.

43. Eberlein CA, Stetson D, Markovets AA, Al-Kadhimi KJ, Lai Z, Fisher PR, et al. Acquired resistance to the mutant-selective EGFR inhibitor AZD9291 is associated with increased dependence on RAS signaling in preclinical models. Cancer Research. 2015;75(12):2489–2500.

44. Chakrabarti S, Michor F. Pharmacokinetics and drug-interactions determine optimum combination strategies in computational models of cancer evolution; 2017. Cancer Research (in print).

45. Ewens WJ. Mathematical Population Genetics. I. Theoretical Introduction. New York: Springer; 2004.

46. Traulsen A, Hauert C. Stochastic evolutionary game dynamics. In: Schuster HG, editor. Reviews of Nonlinear Dynamics and Complexity. vol. II. Weinheim: Wiley-VCH; 2009. p. 25–61.

47. Altrock PM, Gokhale CS, Traulsen A. Stochastic slowdown in evolutionary processes. Physical Review E. 2010;82:011925.

48. Altrock PM, Traulsen A, Galla T. The mechanics of stochastic slowdown in evolutionary games. Journal of Theoretical Biology. 2012;311:94–106.

49. Ashcroft P, Altrock PM, Galla T. Fixation in finite populations evolving in fluctuating environments. Journal of the Royal Society Interface. 2014;11:20140663.

50. Ashcroft P, Traulsen A, Galla T. When the mean is not enough: Calculating fixation time distributions in birth-death processes. Physical Review E. 2015;92(4):042154.

51. Wu B, Traulsen A, Gokhale CS. Dynamic properties of evolutionary multi-player games in finite populations. Games. 2013;4(2):182–199.

52. Altrock PM, Traulsen A, Nowak MA. Evolutionary games on cycles with strong selection. Physical Review E. 2017;95:022407.

53. Altrock PM, Traulsen A, Reed FA. Stability properties of underdominance in finite subdivided populations. PLoS Computational Biology. 2011;7:e1002260.

54. Mafessoni F, Lachmann M. Selective strolls: fixation and extinction in diploids are slower for weakly selected mutations than for neutral ones. Genetics. 2015;201(4):1581–1589.

55. Hilbe C, Abou Chakra M, Altrock PM, Traulsen A. The evolution of strategic timing in collective-risk dilemmas. PLoS One. 2013;6:e66490.

56. Abou Chakra M, Traulsen A. Evolutionary dynamics of strategic behavior in a collective-risk dilemma. PLoS Computational Biology. 2012;8:e1002652.

57. Yeates JA, Hilbe C, Zwick M, Nowak MA, Lehman N. Dynamics of prebiotic RNA reproduction illuminated by chemical game theory. Proceedings of the National Academy of Sciences. 2016;113(18):5030–5035.

58. Anderson AR, Hassanein M, Branch KM, Lu J, Lobdell NA, Maier J, et al. Microenvironmental independence associated with tumor progression. Cancer Research. 2009;69:8797–8806.

59. Basanta D, Scott JG, Rockne R, Swanson KR, Anderson AR. The role of IDH1 mutated tumour cells in secondary glioblastomas: an evolutionary game theoretical view. Physical Biology. 2011;8(1):015016. 20

60. Archetti M, Scheuring I, Hoffman M, Frederickson ME, Pierce NE, Yu DW. Economic game theory for mutualism and cooperation. Ecology Letters. 2011;14(12):1300–1312.

61. Kaznatcheev A, Scott JG, Basanta D. Edge effects in game-theoretic dynamics of spatially structured tumours. Journal of The Royal Society Interface. 2015;12(108):20150154.

62. Li XY, Pietschke C, Fraune S, Altrock PM, Bosch TCG, Traulsen A. Which evolutionary games do growing bacterial populations play? Journal of the Royal Society Interface. 2015;12:20150121.

63. Bozic I, Reiter JG, Allen B, Antal T, Chatterjee K, Shah P, et al. Evolutionary dynamics of cancer in response to targeted combination therapy. Elife. 2013;2.

64. Werner B, Scott JG, Sottoriva A, Anderson ARA, Traulsen A, Altrock PM. The cancer stem cell fraction in hierarchically organized tumors can be estimated using mathematical modeling and patientspecific treatment trajectories. Cancer Research. 2016;76:1705–1713.

65. Tang M, Zhao R, van de Velde H, Tross JG, Mitsiades C, Viselli S, et al. Myeloma cell dynamics in response to treatment supports a model of hierarchical differentiation and clonal evolution. Clinical Cancer Research. 2016;22(16):4206–4214.

66. Ibrahim-Hashim A, Robertson-Tessi M, Enriquez-Navas PM, Damaghi M, Balagurunathan Y, Wojtkowiak JW, et al. Defining cancer subpopulations by adaptive strategies rather than molecular properties provides novel insights into intratumoral evolution. Cancer Research. 2017;77(9):2242–2254.

67. Kaznatcheev A, Peacock J, Basanta D, Marusyk A, Scott JG. Cancer associated fibroblasts and alectinib switch the evolutionary games that non-small cell lung cancer plays. bioRxiv. 2017;Available from: http://www.biorxiv.org/content/early/2017/08/21/179259.

68. Werner B, Beier F, Hummel S, Balabanov S, Lassay L, Orlikowsky T, et al. Reconstructing the in vivo dynamics of hematopoietic stem cells from telomere length distributions. Elife. 2015;p. e08687v2.

69. Altrock PM, Brendel C, Renella R, Orkin SH, Williams DA, Michor F. Mathematical modeling of erythrocyte chimerism informs genetic intervention strategies for sickle cell disease. American Journal of Hematology. 2016;91(9):931–937.

